# Non-parametric test for connectivity detection in multivariate autoregressive networks and application to multiunit activity data

**DOI:** 10.1101/100669

**Authors:** M Gilson, A Tauste Campo, X Chen, A Thiele, G Deco

## Abstract

Directed connectivity inference has become a cornerstone in neuroscience to analyze multivariate data from neuroimaging and electrophysiological techniques. Here we propose a non-parametric significance method to test the non-zero values of multivariate autoregressive model to infer interactions in recurrent networks. We use random permutations or circular shifts of the original time series to generate the null-hypothesis distributions. The underlying network model is the same as used in multivariate Granger causality, but our test relies on the autoregressive coefficients instead of error residuals. By means of numerical simulation over multiple network configurations, we show that this method achieves a good control of false positives - type 1 error - and detects existing pairwise connections more accurately than using the standard parametric test for the ratio of error residuals. In practice, our method aims to detect temporal interactions in real neuronal networks with nodes possibly exhibiting redundant activity. As a proof of concept, we apply our method to multiunit activity (MUA) recorded from Utah electrode arrays in a monkey and examine detected interactions between 25 channels. We show that during stimulus presentation our method detects a large number of interactions that cannot be solely explained by the increase in the MUA level.

## INTRODUCTION

In recent years, there has been a growing interest in developing multivariate techniques to infer causal relations among time series. The initial formulation of the problem goes back to the seminal work by Granger in 1960’s (Granger, 1969) motivated by the analysis of the pairwise influence between economic time series. In this work, Granger decomposes the cross-spectrum of two autoregressive time series into two directional components that account for the potential causal influences between each other. A general solution of the problem in multivariate scenarios was developed a decade later by the introduction of multivariate autoregressive (MVAR) processes, which allow the estimation of causal relationships between nodes in networks with linear feedback based on their observed activity (Amemiya, 1974; Geweke, 1982, 1984; Lütkepohl, 2005). The MVAR was further combined with spectral analysis to develop the directed transfer entropy function (Kamiński & Blinowska, 1991; Kamiñski, Ding, Truccolo, & Bressler, 2001), which has been employed to analyze connectivity patterns in neurobiological systems (Babiloni et al., 2005; Wilke, Ding, & He, 2008). Granger causality analysis is nowadays often used to evaluate the influence of a group of variables onto other, which corresponds to the influence of a subgroup of nodes onto another one in networks. Nevertheless, it it also applied to detect individual connections between pairs of nodes (each subgroup being a single node), which sets the context of the present paper.

In neuroscience, this inference problem has been transposed to analyze interactions between neuronal populations from spiking activity or neuroimaging measurements such as fMRI, EEG and MEG (Lusch, Maia, & Kutz, 2016; Messé, Rudrauf, Benali, & Marrelec, 2014; Michalareas, Schoffelen, Paterson, & Gross, 2013; Rogers, Katwal, Morgan, Asplund, & Gore, 2010; Seth, Barrett, & Barnett, 2015; Storkey et al., 2007). Two types of estimation procedures may be distinguished: measures relying on an underlying interaction model such as Granger causality analysis (M. Ding, Chen, & Bressler, 2006) and dynamic causal modeling (DCM) (Friston, Harrison, & Penny, 2003) on the one hand; and model-free measures such as transfer entropy (Schreiber, 2000) and directed information (Massey, 1990), which make minimal model assumptions on the other hand. Although model-free approaches have proven useful to describe neural propagation at spike-train level (So, Koralek, Ganguly, Gastpar, & Carmena, 2012; Tauste Campo et al., 2015), certain assumptions are required when estimating interactions at the neuronal population level, in which broader spatial and temporal scales contribute to shaping the signals. Motivated by data-driven practical problems, methodological refinements of Granger causality analysis (or MVAR-based methods) have considered additive noise (Vinck et al., 2015) or measurement noise via state-space models (Barnett & Seth, 2015; Friston et al., 2014). However, in the majority of cases, the ratio behind the detection test concerns sub-model error residuals, which might become too similar when connections are placed in a highly redundant network, thus increasing the missed detection rate (Stramaglia, Cortes, & Marinazzo, 2014).

To overcome the difficulties of detecting directed connections in the general context of large networks, we propose to test the significance of the MVAR coefficients using a non-parametric procedure. As a generative model, the MVAR process is canonically related to Granger causality analysis: the linear regression in the upper right inset of Fig. 1A provides both coefficients and residuals, the latter being viewed as the remaining uncertainty in the prediction of the target time series by its source(s). By comparing the residuals of two linear regressions - one involving a supposed driver node and one without it - in a log ratio, traditional tests for Granger causality estimate the effective interaction of one node onto another (Barrett & Barnett, 2013). Since these log ratios asymptotically converge to known distributions, parametric statistical tests have been developed to assess the significance of these interactions (Barnett & Seth, 2014). Instead, our proposed method evaluates the significance of the MVAR coefficients to infer the existence of network connections. To achieve this, we propose a non-parametric significance test in the regression coefficients space. Previous literature on non-parametric testing for Granger causality has resorted to surrogate data generated by trial shuffling (Dhamala, Rangarajan, & Ding, 2008; Nedungadi, Rangarajan, Jain, & Ding, 2009), bootstrap procedures (Diks & DeGoede, 2001) or by phase randomization in frequency-domain measures (L. Ding, Worrell, Lagerlund, & He, 2007; Li et al., 2016). Here we focus on within-trial surrogate tests for time-domain coefficients and compare them across standard techniques (Faes, Marinazzo, Montalto, & Nollo, 2014; Schreiber & Schmitz, 1996; Winkler, Ridgway, Webster, Smith, & Nichols, 2014).

**Figure 1.**
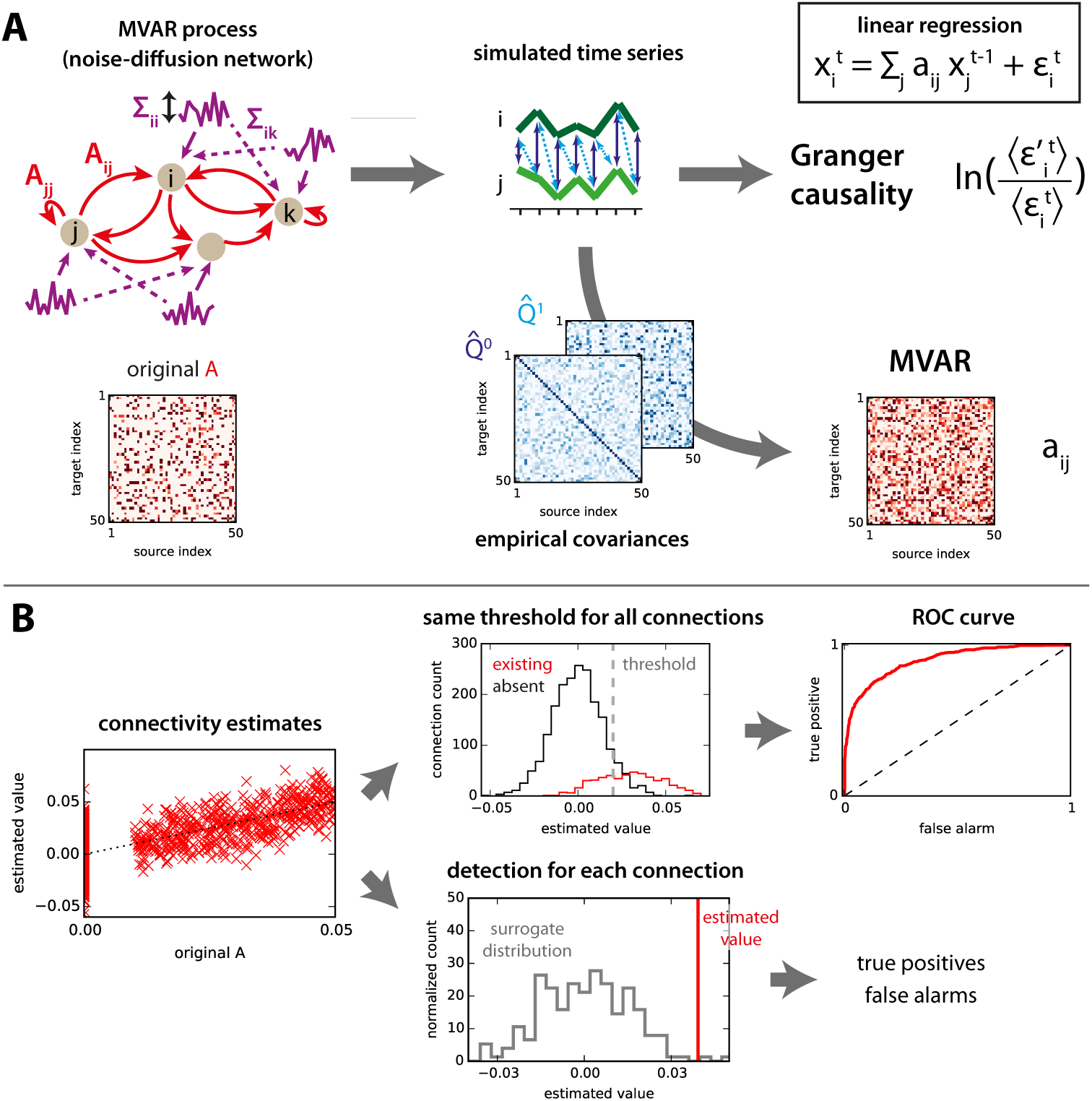
Network model and connectivity estimation. **A:** For a given directed connectivity *A* and input covariances Σ (left), the network activity (middle) is simulated using Eq. (1). From the observed time series, the existing interactions in the original connectivity can be estimated (right): Granger causality analysis uses the residuals of linear regressions (∈ in the upper right equation; see Methods for details about the residuals used in the log ratio), whereas MVAR corresponds to the coefficients. Note that MVAR can be obtained using the empirical covariance matrices 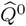 and 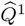, see in Eq. (7) for *τ* = 0 and 1. **B:** The left panel compares the estimated values to the original values for all connections in the network. The upper thread displays the distributions of estimated values for existing and non-existing connections in the original network. Using a sliding threshold (vertical dashed gray line) on the estimated values, one can calculate the ROC curve (right). The lower thread compares the estimated value for a single connection to a null distribution. From this, the choice whether the connection exists or not is made for each individual connection, yielding a single pair of true-positive and false-alarm rates.

Our approach is motivated by the growing of multichannel recording techniques in neuroscience, which require tailored multivariate analysis. In the context of recurrent networks, which are ubiquitous in neuroscience, we provide numerical evidence that these tests can achieve a good control of the false-alarm rate and might improve the miss rate by properly adapting the null distribution to each connection. The focus of the present analysis is on the case where we observe more time samples (a few thousands per node) than the network size (about a hundred nodes), a usual ground for electrophysiological data. Within this regime, we test the robustness of the detection method for a broad range of network parameters and various non-trivial topologies inspired by neuronal networks.

## METHODS: MULTIVARIATE AUTOREGRESSIVE MODEL AND CONNECTIVITY ESTIMATION

The activity in the MVAR process - a.k.a. noise-diffusion discrete-time network - is described by the following equation:
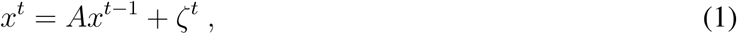

where the connection matrix *A* describes the interactions between coordinates of the vector 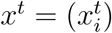 with time *t* being an integer and node index 1 ≤ *i* ≤ *N*. Here we constrain our study to the case where *ζ^t^* is Gaussian (possibly cross-correlated noise), whose realizations are time independent for successive time steps. Without loss of generality, we assume that all variables *ζ^t^* have zero means, giving the same property for all *x_i_*. We only consider MVAR processes of order 1 in a first place, but will extend the work to the case of order 2 in a later section.

### Granger causality analysis

Granger causality analysis is usually presented using time series and the estimation of non-zero coefficients in A from observed activity over a period 1 ≤ *t* ≤ *T* relies on the linear regression of the activity 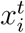 of a given node *i* at time *t* by the past activity of a subset *S* of network nodes:
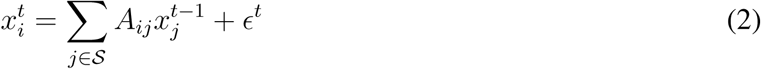

for 2 ≤ *t* ≤ *T*. When *T* is large, the coefficients *a_ij_* converge toward *A_ij_*. We define the residual *ϵ* as the standard deviation of the *ϵ^t^* for the ordinary least-square (OLS) regression in Eq. (2), which is for
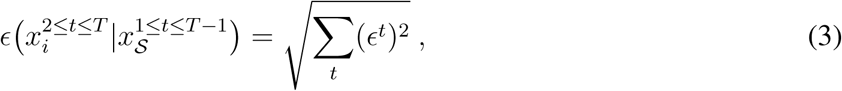

with a notation similar to conditional probabilities; the superscript *t* indicates the considered time range and the subscripts indicate the nodes involved. To detect the existence of connection *j* → *i* in a network, two types of Granger causality analysis exist: ‘unconditional’ and ‘conditional’ (Geweke, 1982, 1984).,
they consider the comparisons of the following residuals:
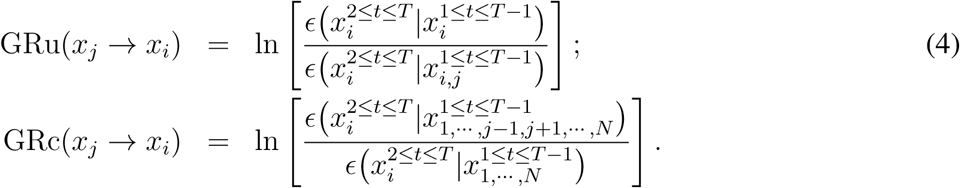

For both GRu and GRc, which have a univariate target node *x_i_*, the usual parametric test for significance relies on the F statistics, which performs better for small number of samples (Barnett & Seth, 2014). The null hypothesis of no interaction for GRu(*x_j_* → *x_i_*) corresponds to *m* = *T*, *p* = 1, *n_x_* = 1 and *n_y_* = 2 using the notation in Barnett and Seth (2014)
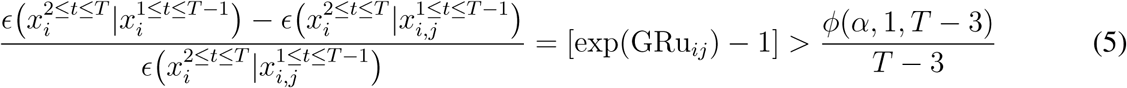

with *α* the desired sensitivity and *ϕ* the inverse survival function of the F-distribution (<u>www.scipy.org</u>, n.d.). The equivalent for GRc corresponds to *n_y_* = *N*, yielding
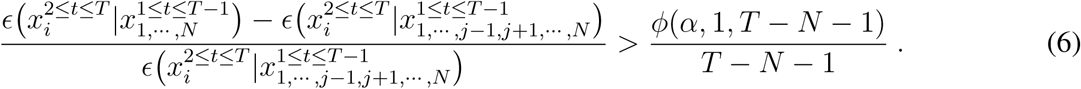

We also use non-parametric tests for GRc by performing a circular shift (see details below in Section ‘Generation of surrogate time series’) either on the target node for each connection (Faes et al., 2014) or independently on the time series of all nodes (in order to save time in estimating the full network’s connectivity by shuffling somehow all targets simultaneously). Both cases provide a null distribution for the log ratio, with which the actual estimated log ratio can be compared.

### Multivariate autoregressive (MVAR) estimation

To detect the existence of connections *A_ij_* > 0, another possibility is to estimate the coefficients themselves, which can be done using the covariances of the observed activity variables *x^t^* (Lütkepohl, 2005):
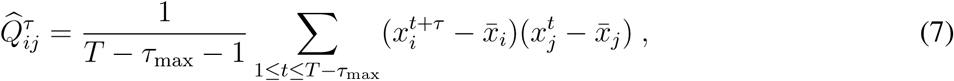

where *T* denotes the number of successive samples indexed by *t, τ* ∈ {0, 1} is the time shift (here *τ_max_* = 1) and the observed mean activity for each node is 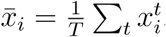. The Yule-Walker equation gives a consistency equation for the theoretical covariance matrices (without hat) in terms of the connectivity A in the dynamics described by Eq.(1):
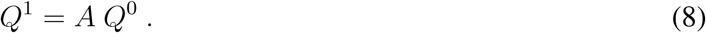

The estimation of network connections relies on evaluating A from Eq. (8) for the empirical covariance matrices defined as Eq. (7) and calculated for a given time series:
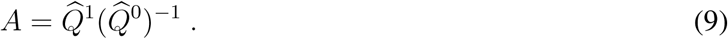

Note that this OLS estimate corresponds to the linear regression related to 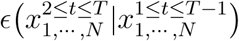 and is also to the linear model with maximum likelihood under the assumption that the observed process is Gaussian.

### MVAR of order 2

Eq. (1) can be extended to the case where the activity vector *x*^*t*^ is determined by the two previous time steps:
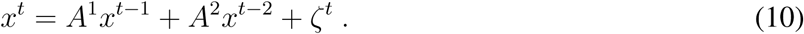

For the second order, we use *τ_max_* = 2 in Eq. (7) and the estimation of *A*^1^ and *A*^2^ via the Yule -Walker equation is given by (Lütkepohl, 2005, p. 86)
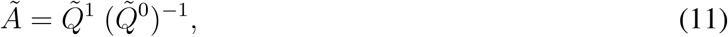

with the block matrices
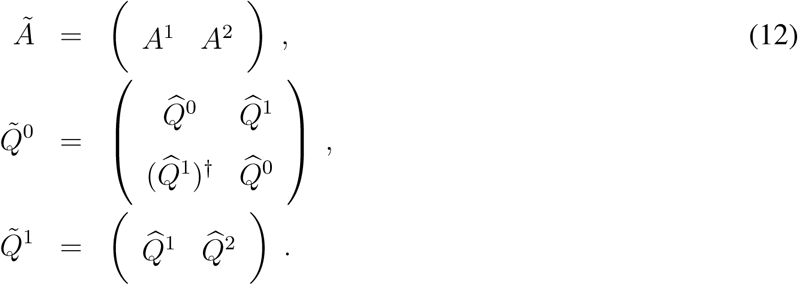

The coefficients of *A*^1^ and *A*^2^ can thus be estimated using a matrix multiplication and an inversion involving the covariances, as with the first-order case in Eq. (9).

### Generation of surrogate time series

In this paper, we consider circular shifts (CS), random permutations (RP) and phase randomization (PR) to shuffle the time points of the observed time series. From the original 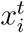 with 1 ≤ *t* ≤ *T*,

CS draws a random integer 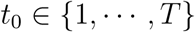 and returns 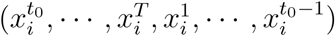;
RP draws a random permutation *σ* of 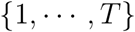 such that each integer appears once (and only once) and returns 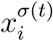;
PR calculates the discrete Fourier transform 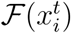 of the original 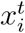, then multiplies each of the *T* coefficients of 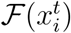 by exp(2*πιϕ^t^*) with *ϕ^t^* randomly chosen in [0, 2*π*], and performs the inverse transform.

Importantly, these operations are applied to each time series independently of the others.

In addition, we consider the replacement of all time series in the network by *T* normally distributed variables with a standard deviation equal to the mean of the standard deviations of 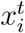 along the time axis, then averaged for all nodes. We refer to these surrogates as STD.

### Experimental setup and processing of electrode measurements to extract MUAe activity

All procedures were carried out in accordance with the European Communities Council Directive RL 2010/63/EC, the US National Institutes of Health Guidelines for the Care and Use of Animals for Experimental Procedures, and the UK Animals Scientific Procedures Act. Two male macaque monkeys (5 - 14 years of age) were used in the experiment; only the data for the first one is used here. A surgical operation was performed under sterile conditions, in which a custom-made head post (Peek, Tecapeek) was embedded into a dental acrylic head stage. Details of surgical procedures and post-operative care have been published previously (Thiele, Delicato, Roberts, & Gieselmann, 2006). During the surgery microelectrode chronic Utah arrays (5*5 grids), attached to a CerePort base (Blackrock Microsystems) were implanted into V1. Electrodes were 1 mm in length in line with procedures described in Supèr and Roelfsema (2005). Stimulus presentation was controlled using CORTEX software (Laboratory of Neuropsychology, NIMH, http://dally.nimh.nih.gov/index.html) on a computer with an Intel Core i3-540 processor. Stimuli were displayed at a viewing distance of 0.54 m, on a 25” Sony Trinitron CRT monitor with a resolution of 1280 by 1024 pixels, yielding a resolution of 31.5 pixels / degree of visual angle (dva). The monitor refresh rate was 85 Hz for monkey 1, and 75 Hz for monkey 2. A gamma correction was used to linearize the monitor output, and the gratings had 50% contrast. Monkeys performed a passive viewing task where they fixated centrally while stationary sinusoidal grating of either horizontal or vertical orientation and 2 cycle per degree spatial frequency, were presented in a location that covered all receptive fields recorded from the 25 electrode tips. Stimuli were presented 500 ms after fixation onset for 150 ms. Raw data were acquired at a sampling frequency of 32556 Hz using a 64-channel Digital Lynx 16SX Data Acquisition System (Neuralynx, Inc.). Following each recording session, the raw data were processed offline using commercial (Neuralynx, Inc.). Signals were extracted using Cheetah 5 Data Acquisition Software, with bandpass filtering set to allow for spike extraction (600-9000 Hz) and saved at 16-bit resolution.

In the present study, we focus on the period starting 200 ms before and finishing 200 ms after the stimulus onset, for 4 conditions (vertical gradings with pre/post cue in the receptor/opposite field) that will not be compared in details. The electrode recordings is firstly down-sampled from 32556 Hz to 1000 Hz. A high-pass filter above 400 Hz is then applied - 3rd-order Butterworth filter at 0.8 of the Nyquist frequency (<u>www.scipy.org</u>, n.d.) - followed by a smoothing of 4 ms to extract the envelope of the resulting signal, by retaining the 250 time points of 1000-ms period surrounding the stimulus onset.

## BENCHMARK OF DETECTION PERFORMANCE FOR SYNTHETIC DATA

The workflow of the benchmark for the estimation procedure is schematically represented in Fig. 1A. We first consider a MVAR process defined by Eq. (1) with given connectivity matrix *A* and input covariances (obtained by mixing independent Gaussian processes) to generate the activity of the network. From the observed activity over a period of duration *T*, we estimate the coefficients matrix *A* using the covariances as described in Eq. (9). We also perform the linear regressions of each node activity over the past activity of given subsets of nodes corresponding to the unconditional (GRu) and conditional (GRc) Granger causality analysis, from which we calculate the ratios of residuals in Eq. (4). Actually, these estimates correspond to the same ordinary least-square (OLS) regression (top right in Fig. 1A) and the difference resides in the spaces where they lie: coefficients versus residuals.

For each method, the prediction power can be measured by the relationship between the estimated values and the original connectivity values, as illustrated in Fig. 1B (left). To discriminate between existing and absent connections, one can apply a common threshold for all connections (top thread); by sliding this threshold, we obtain the ROC curve with the rate of false alarms on the x-axis and true positives on the y-axis. The area under the curve indicates in a single value how well the ranking of estimated value performs for the detection of connections in the original connectivity. Alternatively, an individual test can be made for each connection in the network, for example by comparing the estimated value to a null distribution (bottom thread). Here again, we obtain two rates of false positives and negatives.

### Coefficients from linear regression potentially predict better existing connections than residuals

We start with the comparison between the predictability of coefficients and residuals for MV, GRu and GRc for all connections in a given network. To do so, we simulate 500 randomly connected networks, which are simulated with different sizes (*N* = 50 to 150 nodes), density and connectivity weights (uniformly drawn in a randomly chosen range [*w*_min_, *w*_max_]); here inputs are *not* correlated: the *ζ_i_* are independent across node indices in Eq. (1). For each network configuration, we evaluate the accuracy for connection detection via the area under the ROC curve (see the upper thread in Fig. 1B). Fig. 2A displays this ROC-based accuracy as a function of the number of observed time samples (x-axis) represented by violin plots for 500 randomly connected networks. When considering many samples (10^4^), all methods perform well. However, for smaller sample sets, the MVAR method exhibits superior performance than GRu: as measured by the Mann-Whittney test, *p* < 10^−45^, *p* < 10^−19^ and *p* < 10^−5^ for the three values of observed samples, respectively.

**Figure 2.**
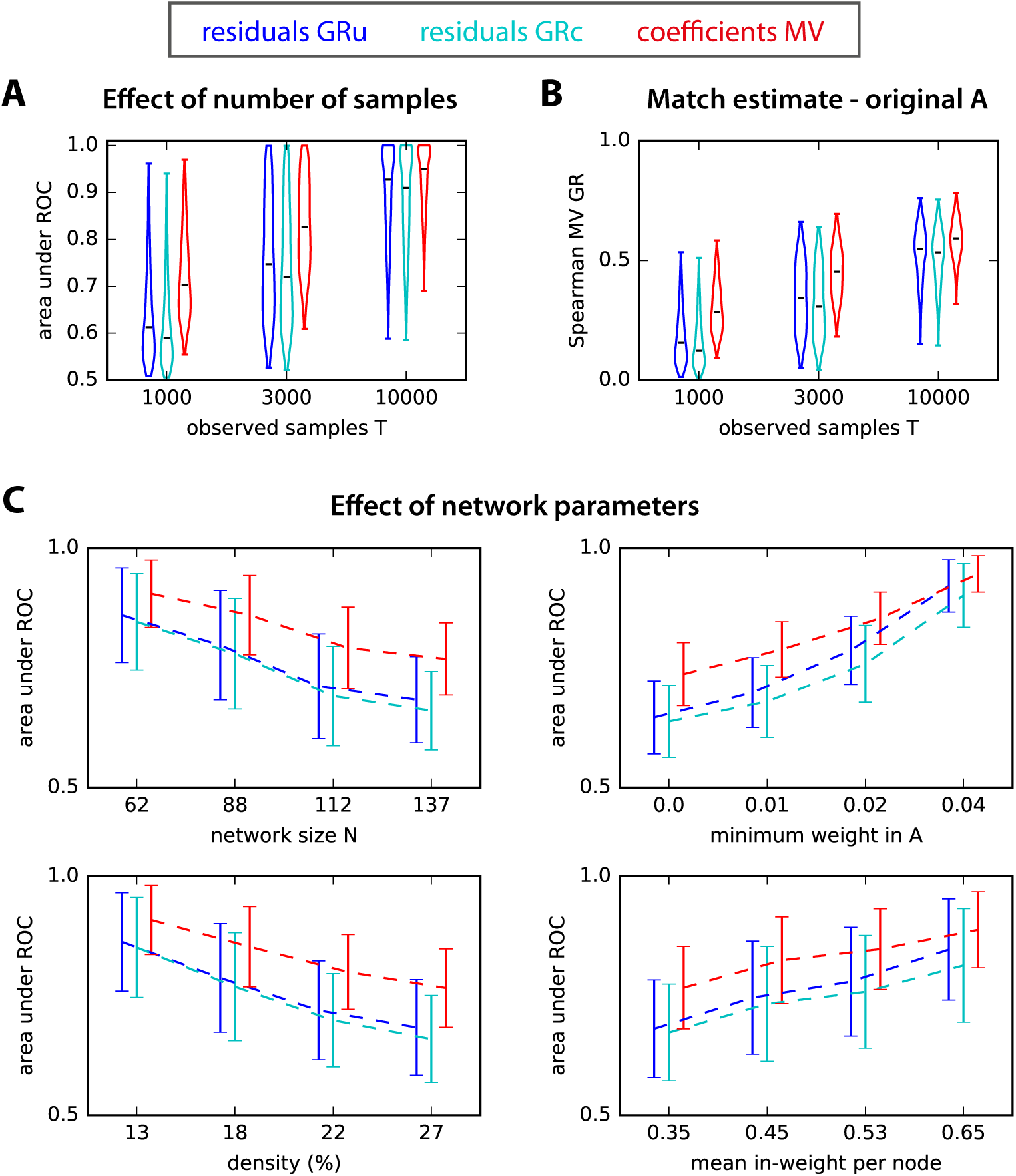
ROC-based prediction power. **A:** Area under ROC for estimated *A* obtain from log-ratios of residuals obtained from Granger causality analysis (unconditional for GRu and conditional for GRc) and MVAR. The x-axis indicates three sample size *T* for the observed network activity. The the violin plots correspond to 500 simulated networks of various sizes and connectivity strengths (the horizontal black bar indicates the median). **B:** Match of the ranking between GRu, GRc and MVAR estimates and the original connectivity weights *A*, as measured by the Spearman correlation coefficient. The plotted values correspond to the 500 networks in A and the x-axis indicates the sample size *T*. **C:** Effect of network parameters on ROC-based performance. Influence of network size *N*, connectivity density, sum of recurrent connectivity strengths, minimum weight *w*_min_ in *A*, mean noise on the diagonal of ∑ and mean off-diagonal noise in ∑ on the ROC-based accuracy in Fig. 2A. In each plot, the network configurations have been grouped in quartiles according to the parameter plotted on the x-axis, and the corresponding group mean and standard deviations are indicated; the curves are displaced horizontally to improve legibility.

Although error residuals ratios are in a different space from the true weights in *A*, one expects some degree of correlation between them, such that Granger causality analysis effectively detects connections. In Fig. 2B, both GRu and GRc estimates have a ranking similar to the original A weights (as measured by the Spearman correlation) for *T* = 10000 observed time samples, but this weakens dramatically for *T* ≤ 3000. In contrast, the ranking for estimated MVAR coefficients reflects much better the original *A* for *T* ≤ 3000. In the studied networks, GRu performs slightly better than GRc. As analyzed in previous studies, this can be consequence of the balance between redundant and synergistic activity exhibited by the simulated network nodes (Stramaglia et al., 2014). To shed light into the effect of the network structure, we next examine how the ROC-based performance in Fig. 2A depends on the controlled network parameters. The four panels in Fig. 2C display the trends of the values for the 500 networks as a function of the network size *N*, the network density, the minimum weight in the original network (*w*_min_ mentioned above) and the mean sum of incoming weights per node. For illustration purpose, the 500 networks are grouped in quartiles for each parameter. Not surprisingly, the estimation accuracy of all methods decreases as a function of the network size *N* and density, and increases as a function of the minimum connectivity weight and the mean incoming weight per node. More interestingly, in challenging configurations with small weights, MVAR consistently shows a superior performance by a larger gap compared to Granger causality analyses. These findings support the use of coefficients to robustly detect connections in recurrently connected networks. Note that GRu performs on average slightly better than GRc here: the discrepancy decreases as a function of the network density, which may follow from lower redundancy in the recurrent network (Stramaglia et al., 2014).

### Arobust non-parametric significance test for MVAR

We have so far examined the performance of different estimation methods based on the area under the ROC curve, which corresponds to a single threshold for all connections in a network and combines the information about false alarms and true detection over the whole range of estimated values. However, in the context of real data, the decision for the existence of a connection typically relies on comparing the value of the connection estimate with a given statistical threshold. For GRu and GRc, such parametric tests have been developed, for example, based on the F statistics (Barnett & Seth, 2014). Equivalently, it is sufficient to know how the values of the estimates for absent connections are distributed, in order to select a desired rate of false alarms (type-1 error). In this section, we develop a significance test for the estimated MVAR coefficients by providing a null-hypothesis distribution for absent connections.

Our approach relies on the fact that covariances reflect the underlying connectivity: we thus construct the null distribution for estimates by performing a random permutation for each of the observed time samples, which “destroys” the covariance structure apart from the variances on the diagonal of 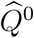 as illustrated in Fig. 3A; other methods will be tested in a later section. From the resulting covariance matrices, we evaluate a surrogate connectivity matrix. The core result underlying our surrogate approach is shown in Fig. 3B: the distribution of surrogate estimates (thick black line) is compared against the distribution of existing (red) and absent (blue) connections in a simulated random network model: the surrogate distribution in black provides a good approximation for the distribution of estimates for non-existing connections in blue.

**Figure 3.**
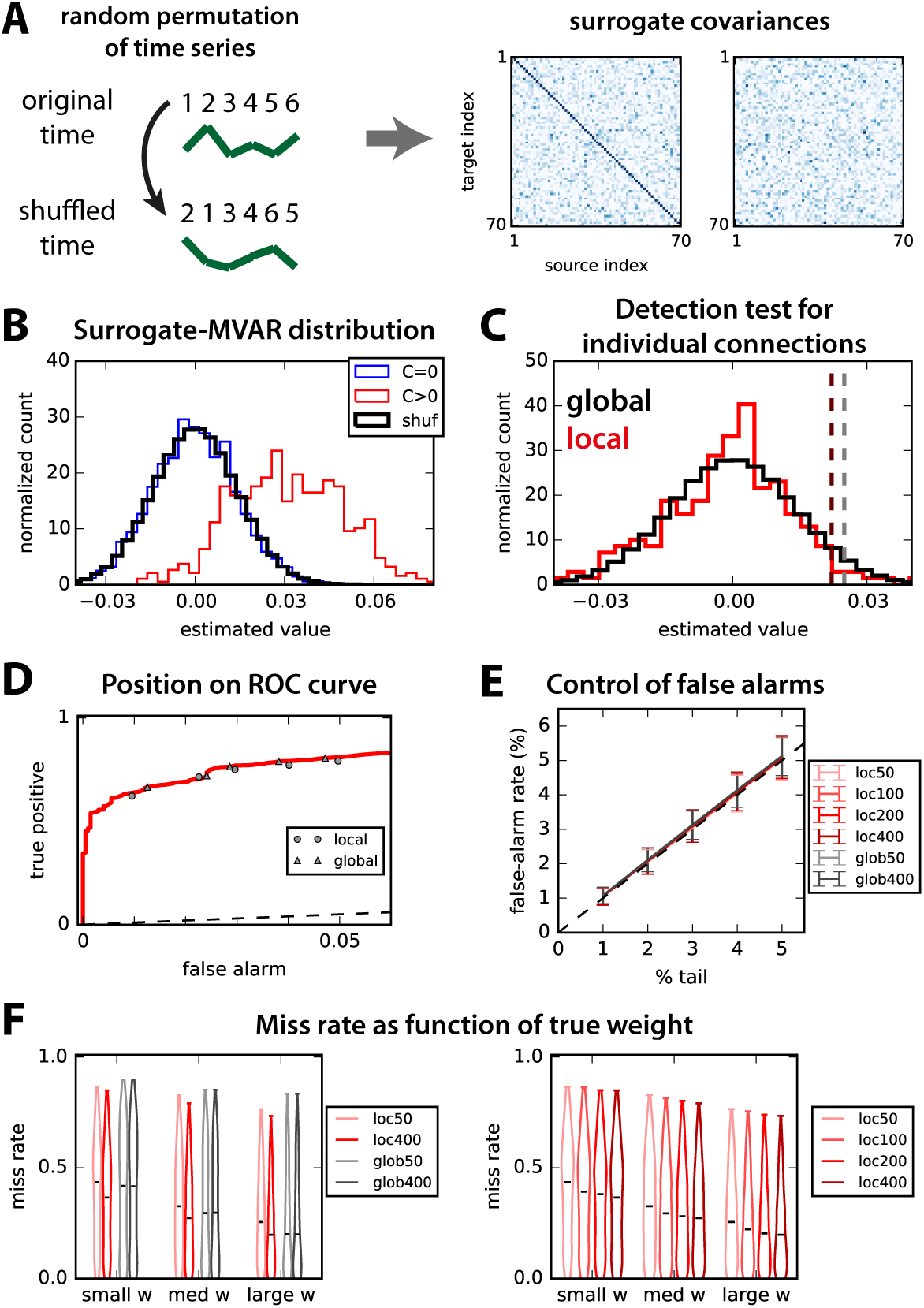
Non-parametric test to assess significance for MVAR coefficients based on random permutations. **A:** Schematic illustration of random permutation (numbers indicate time) applied independently to all observed time series (green curves) to generate the surrogate covariances (right panels). **B:** Pooled distributions of estimated weights for the existing (in red) and absent (blue) connections. The thick black curve indicates the distribution of connections over 100 surrogates, which closely matches the blue distribution. **C:** For a given connection, we compare two methods: for ‘local’ in red, the null distribution corresponds to the matrix elements for the same connection in 200 surrogates; for ‘global’ in black, the null distribution is the pooled distribution for all *N*^2^ elements of the 200 surrogate matrices (same as in B). The dark red and gray dashed lines indicate the detection thresholds corresponding to the 4% tail for those two options. **D:** The performance of the two non-parametric methods for the thresholds described in B and C is displayed on the ROC curve for a desired false-alarm rate ranging from 1 to 5%. Triangles indicate the local test and circles the global test. **E:** Comparison of the desired (% of the tail of null distribution) and actual rate of false alarms for the local and global tests when varying the the number *S* of surrogates (see figure in legend). Error bars indicate one standard deviation for 500 random networks; importantly, inputs for these networks have cross-correlation, unlike Fig. 2. **F:** Influence of the strength of original weight on the detection performance for the 500 random networks and a desired false-alarm rate set to 2% in E. In both panels, lighter colors indicate smaller numbers of surrogates *S*, in red for the local test and gray for the global test (see legends).

We consider two options - corresponding to the two threads in Fig. 1B - to test the existence of a connection from an MVAR estimate while keeping the false alarm rate under control.

- The *global* test relies on the null distribution corresponding to the black histogram in Fig. 3C, which is obtained by grouping together all *SN*^2^ matrix elements of all matrices for *S* = 200 surrogates. From that surrogate distribution, we perform a detection test by setting a threshold corresponding to a percentage of the right tail equal to the desired false-alarm rate (here 2%), as illustrated by the vertical gray dashed line.
- Instead, the *local* test uses for each connection the surrogate distribution of *S* values, corresponding to the same matrix element in each of the *S* surrogates. From that distribution in red in Fig. 3C, the detection threshold is defined similarly (vertical dark red dashed line).

The rationale behind these two choices lies in the trade-off between taking into account spatial heterogeneity in the network and gaining larger sample size, as illustrated by the distinct thresholds in Fig. 3C. Note also that the F statistical test for Granger causality analysis corresponds to a global threshold on the log ratio values. When varying the desired false-alarm rate, the two tests perform well, as illustrated in Fig. 3D by their location close to the ROC curve (circles and triangles for local and global, respectively).

To assess the effect of the small variability observed in Fig. 3D over the randomness of network configurations, we simulate 500 randomly connected networks with the same parameters as in Fig. 2, except for the size 50 ≤ *N* ≤ 90 and the presence of input cross-correlations. Note that, from Fig. 2, the chosen size *N* corresponds to a situation where Granger causality analysis performs relatively well as compared to MVAR. The control of the false-alarm rate is displayed in Fig. 3E for both local and global tests with various numbers *S* of surrogates. The control of false-alarm rates is close to perfect across various values for all *S* and both tests (local and global), demonstrating the robustness of the proposed method for randomly connected networks. Following, we fix the desired false-alarm rate to 2% and evaluate the miss rate (true negatives) of both methods depending on the actual weight strength: in Fig. 3F, connections are grouped in terciles for each network configuration. Interestingly, the local test improves with the number *S* of surrogates (right panel), whereas the global test exhibits a constant performance for all *S* (left panel). Note that the advantage of the local test over the global test particularly concerns connections with small weights, which are difficult to detect, in line with Fig. 2C (see influence of the minimum original weight).

Because GRu does not take all network nodes into account, the presence of spatially correlated noise (indicated by the purple dashed arrows in Fig. 1A) dramatically affects the false-alarm rate when using the parametric significance F-test (Barnett & Seth, 2014), as shown in Fig. 4A by the dark blue dashed curve. This is solved by the “complete” linear regression in GRc, achieving a quasi perfect control irrespective of the input correlation level for both the parametric and two non-parametric tests (cyan, green and blue-green dashed curves, respectively), as our non-parametric tests do (red and gray; recall also Fig. 3E). We consider two non-parametric tests for GRc: ‘T’ stands for target, where the null distribution of a connection is obtained by shuffling only the target, and ‘F’ stands for full, where we shuffle all time series simultaneously as in our coefficient-based tests. Both perform equally in terms of false alarms.

The main result of the paper is described in Fig. 4B, where the dashed line corresponds to the miss rate for parametric GRc: our non-parametric method exhibits a better than miss rate - i.e., decrease - for both local and global tests (in red and gray, respectively) on average over the same 500 random networks as in Fig. 4A. For *S* ≥ 200, the local test even becomes better in all cases. Note that the small miss-rate improvements of about 7% actually corresponds to more than 50 existing connections per network here. In contrast, both non-parametric tests for GRc perform worse than the parametric test here, with the target-shuffling surrogate converging faster close to the non-parametric GRc. Fig. 4C displays the trends of the performance of all 5 tests in Fig. 4B as a function of two network properties: the mean incoming weight per node (left) and the density (right). The main result here is that the local test performs better especially in difficult configurations with small weights and dense connectivity.

**Figure 4.**
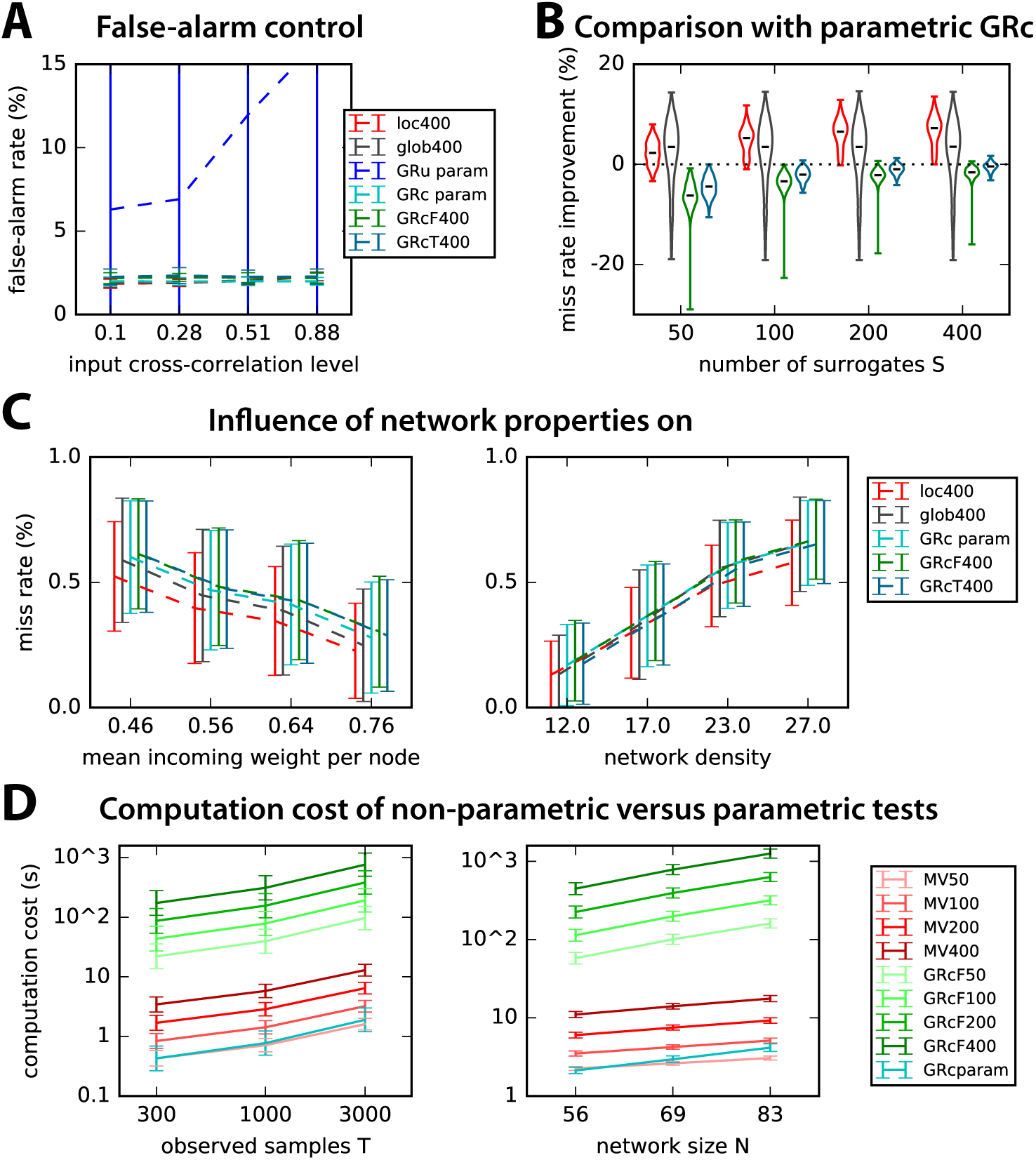
Comparison of our coefficient-based method with Granger causality analysis. **A:** Comparison of the parametric tests for GRu (blue curve) and GRc (cyan) with the non-parametric methods for GRc (green for GRcF and blue-green for GRcT, see the text for details) and MVAR (red for local test and gray for global). The x-axis indicates the strength of input correlations (i.e., pink noise) in the simulated network. The desired false-alarm rate is set to 2% as in Fig. 3F and the number of observed time samples is *T* = 3000. Error bars indicate one standard deviation over the 500 random networks as in Fig. 3E. **B:** Comparison of the miss rate improvement (decrease) with respect to parametric GRc for the 500 networks in A as a function of the number *S* of surrogates (x-axis). Red indicates the local test, gray the global test, green the full-network non-parametric GRc and blue-green the target-only non-parametric GRc. **C:** Details of the performance of the 5 methods in B as a function of the mean incoming weight per node (left) and the network density (right). The plots for the miss rate are similar to those for the ROC-based prediction power in Fig. 2C. **D:** Comparison of the computational cost for the surrogate-based method and parametric tests as a function of the number *T* of observed samples (left) and network size (right). Only GRcF is shown, as GRcT takes much longer time in the unoptimized version that we use.

From Figs. 3F and 4B-C, we conclude that the local test is preferable to the global test provided *S* ≥ 200 surrogates are generated. However, the computational cost increases linearly with *S*, as illustrated in Fig. 4D by the red curves. Note that the parametric GRc (in cyan) takes the same time to calculate as *S* = 50 surrogates. However, our non-parametric method scales better than parametric GRc when the network size increases. As a comparison, the full-network non-parametric test for GRc takes longer time to compute, but further optimization of the calculations could be made that were not incorporated here.

### Comparison of generation methods for surrogates for non-parametric MVAR

The fact that the OLS MVAR estimates can be obtained via the two covariance matrices (with and without time shift, see Fig. 1A) hints at possible methods to generate surrogate by destroying the information in these covariances. Methods to generate surrogate time series have been widely used in the past: circular shifts of the time series (Faes et al., 2014), random permutation (Winkler et al., 2014) and phase randomization (Schreiber & Schmitz, 1996) to generate a null distribution for the ratios in Eq. (4); they are referred to here as CS, RP and PR, respectively. We thus consider these three methods (cf. box in Fig. 5), as well as surrogate time series that only preserve the mean standard deviation averaged over the network (STD), so as test to which extent it is important to preserve the spatial heterogeneity of the nodes’ activity. See Methods for details about the calculations.

**Figure 5.**
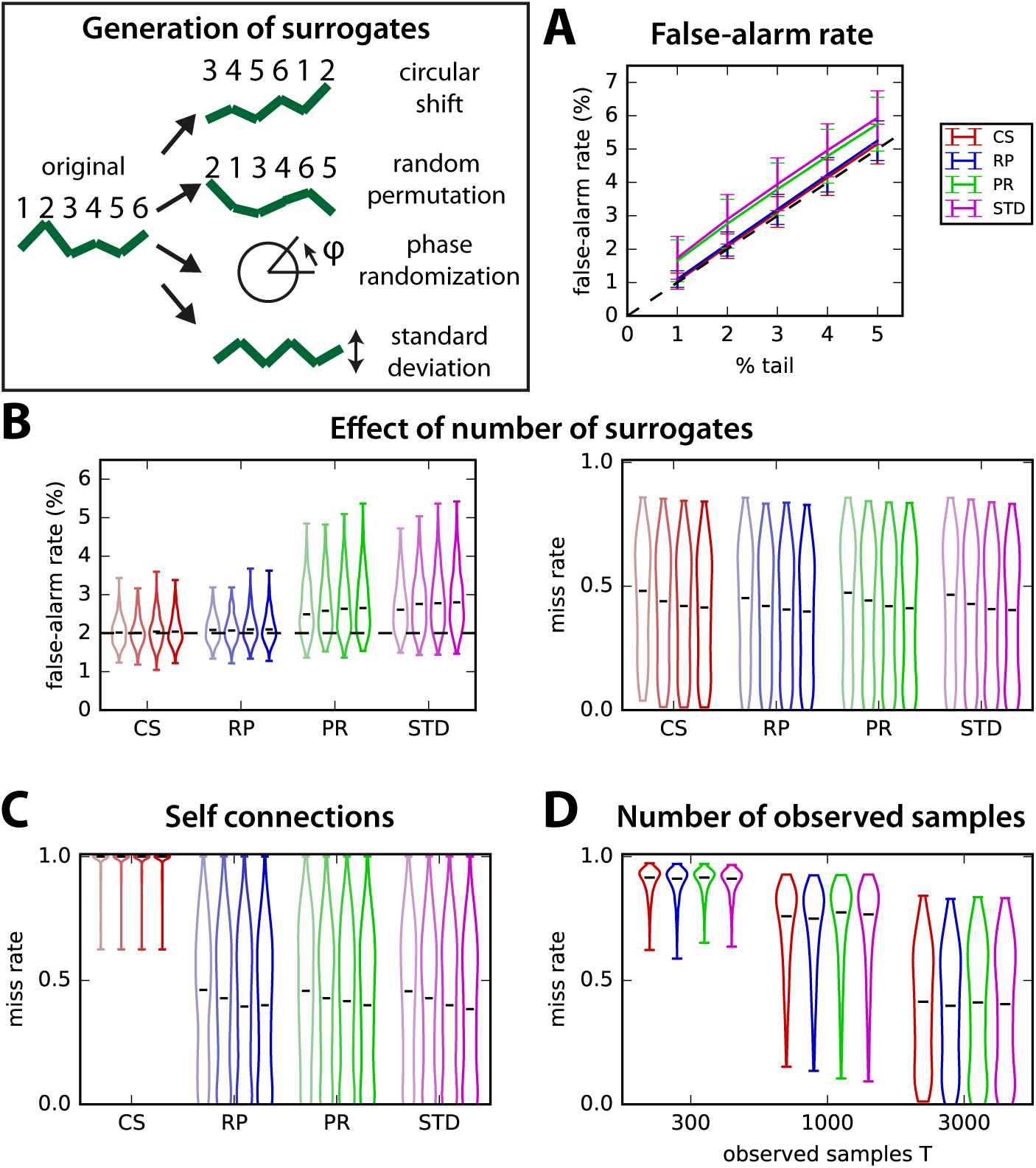
Comparison of the local test for 4 surrogate generation methods. Surrogates are generated by independently performing for each original time series 1) random permutations, 2) circular shifts, 3) phase randomization and 4) replacing the original by a new random time series with the mean standard deviation averaged over all of the original time series in the network. **A:** Comparison of the control of the false-alarm rate for various thresholds on the tail of the distributions of 400 surrogates (% indicated on the x-axis). The error bars correspond to the variability over 500 random networks similar to Fig. 3 with *T* = 3000 observed time samples and the local test. **B:** Influence of the number *S* of surrogates on the detection performance for the 4 surrogate methods (x-axis). The violin plots indicate the distribution of false-alarm rates (left panel) and miss rates (right) for the 500 networks in A with a desired false-alarm rate set to 2% (dashed line in the left panel). Lighter to darker colors correspond to 50, 100, 200 and 400 surrogates, respectively. **C:** Same as the miss rate in B, but only for self connections. **D:** Influence of the number *T* of observed time samples on the miss rate for *S* = 400 surrogates.

The control of false alarms for local tests in Fig. 5A and B is better for CS and RP, whereas the detection of true connections is similar for the four methods over 500 random networks of size *N* = 70. However, CS fails to detect self connections (Fig. 5C). The reason is that, because CS surrogates preserve the autocovariances in the time-shifted covariance, they fail to build a proper null distribution for self-connections. The influence of the number of samples used in the estimation is similar for all methods, as illustrated in Fig. 5D. The comparison with STD (purple), which averages the covariance statistics over the whole network, suggests that the local test makes a good use of the heterogeneous information across nodes. As a conclusion, we retain RP as the best option.

### Influence of network topology

In this part, we test and compare the robustness of global and local surrogate-based detection tests to specific connections and topological configurations. Here, *T* = 3000 observed samples and we compare the local and global tests with *S* = 400 surrogates for 500 networks of each type. In all cases, the simulated networks have the same size *N* = 70, but vary in connectivity density, distribution of recurrent weights and level of input cross-correlation. We compare the miss rate for unidirectional, reciprocal and self connections in the random networks examined until now (and a desired 2% of false alarms). Fig. 6A shows that the miss rate is similar in unidirectional and reciprocal connections with the local test, which performs slightly better than the global test (as in Fig. 3F).

**Figure 6.**
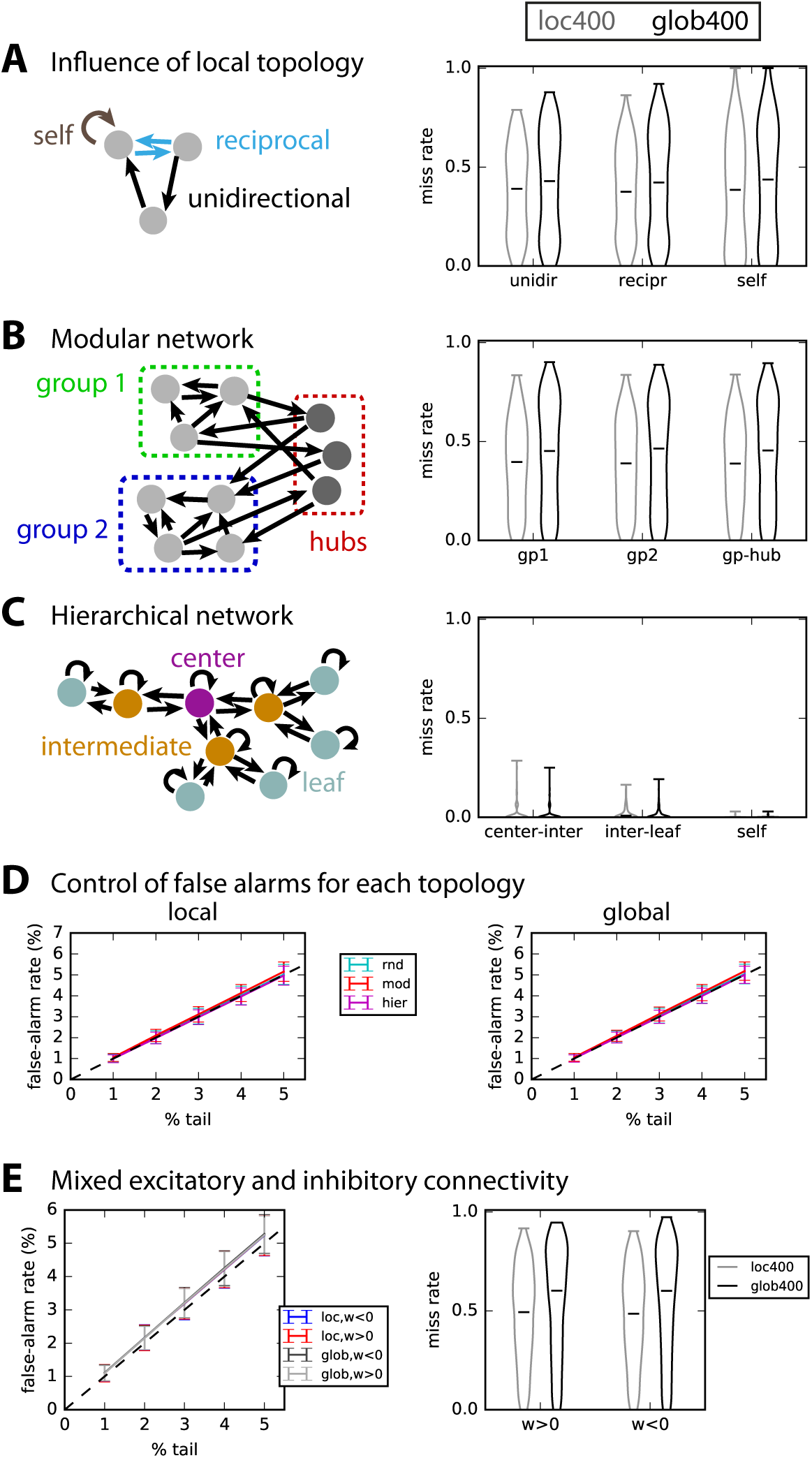
Robustness to non-trivial network topology. **A:** Detection performance for unidirectional, reciprocal and self connections in 500 the randomly connected networks used so far in Fig. 3. **B:** Detection performance for modular topology schematically represented in the left: each of the 500 networks comprises of two groups connected by hubs. The connections are separated depending on whether the connect group nodes or hubs, as indicated by the diagram on the left. **C:** Similar to B with a hierarchical topology, where connections are grouped in 3 subsets: from center to intermediate nodes; from intermediate nodes to leaves; self connections. The results concern 500 such networks, which all have very low density. **D:** Control of false-alarm rate for the local (left) and global (right) significance tests and the three network topologies; the plot is similar to Fig. 3E. **E:** Control of false-alarm rate and miss rate for networks with both excitatory and inhibitory connections.

Now we consider more elaborate network topologies than the random connectivity (Erdös-Rényi) considered so far, namely modular and hierarchical networks. In Fig. 6B, we simulate 500 modular networks with two groups (green and blue) linked by hubs (red, about 5 to 15% of the nodes). Interestingly, intra-group and hub-group connections have a similar miss rate with regard to using local and global surrogates. In Fig. 6C, we simulate hierarchical networks of three layers, for which connections either link the center and an intermediate node, or link an intermediate node and a leaf, or are self connections. This network type is much sparser than the two types in A and B, yielding a quasi perfect detection performance for all types of connections (miss rate < 0.1 in Fig. 6C). In all cases, the local test performs better than the global test. However, the control of false-alarm rate is similar for both tests with all topologies, as can be seen in Fig. 6D.

Finally, we consider a network with both excitatory and inhibitory connections (with a inhibitory ratio equal to 5 to 50% of all) and perform the test by defining a threshold on both tails of the null distributions. As can be seen in Fig. 6E, the positive/negative nature of the connection weights affect neither the false-alarm nor the miss rate. However, the performance is poorer than with excitatory connections only.

We conclude that, in those networks with spatial heterogeneity as with randomly connected networks, the local test with an individual null distribution per connection performs better than the global test. Recall that an improvement of the miss rate by 1% in a network with a density of 20% actually corresponds to *N*^2^0.2/100 ≃ 10 existing connections here, so the plotted improvements concern about 50 connections.

### Applicability to second-order MVAR process

As explained in Methods, an MVAR process whose state depends on the two previous time steps can be estimated with the covariances with time shifts *τ* = 0, 1 and 2; see Eqs. (11) and (12) for details. Here we simply focus on random connectivity for the two corresponding matrices *A*^1^ and *A*^2^, with size *N* that is randomly drawn between 30 and 80; we construct *A*^1^ and *A*^2^ such that a connection *j* → *i* cannot be in both matrices, but at most in one. The existing connections are detected with the non-parametric local test relying on RP surrogates for each matrix separately, as a proof of concept. The control of false alarms in Fig. 7A and the overall detection performance in Fig. 7B suggest that our surrogate method can be extended satisfactorily to higher-order MVAR processes. Note that the improvement by generating more surrogates is rather weak here. Importantly, there is no difference between the detection in *A*^1^ and *A*^2^, as demonstrated in Fig. 7C. Last, the network size worsens the miss rate in Fig. 7D, which affects more dramatically the global test as compared to the local test.

**Figure 7.**
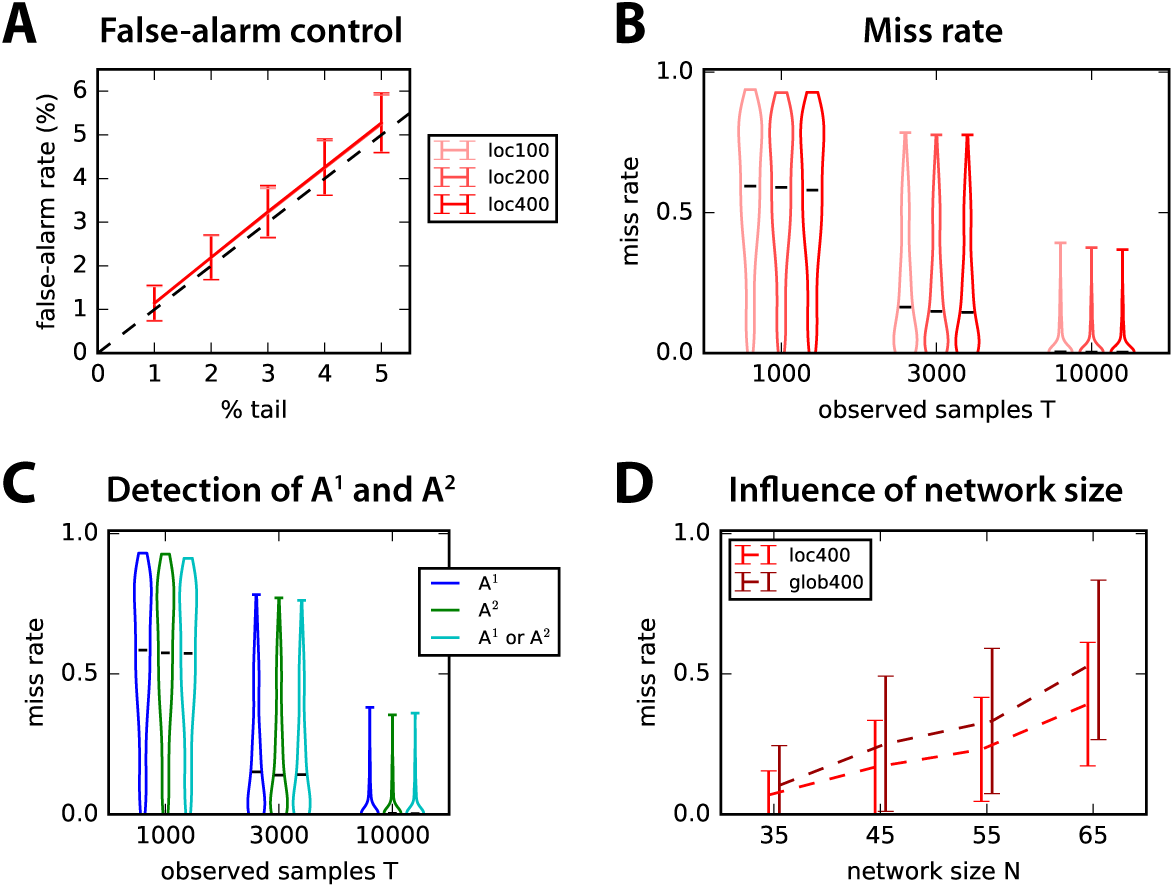
Connectivity detection for second-order MVAR process. **A:** Control of false-alarm rate for the local test with *S* = 100, 200 and 400 in both connectivity matrices *A*^1^ and *A*^2^, corresponding to each time step. Error bars correspond to the variability over 500 network configurations with random connectivity and *T* = 3000 observed samples. **B:** Influence of the number *T* of observed samples (x-axis) on the miss rate for the 500 networks in A. Lighter to darker red indicates the number of surrogates *S*. **C:** Details of the detection performance for *A*^1^ and *A*^2^ separately, as well as connections in either *A*^1^ or *A*^2^. The number of observed samples is indicated on the x-axis as in B. **D:** Influence of the network size *N* on the detection performance in C for the local and global tests with *S* = 400 surrogates.

## APPLICATION TO EXPERIMENTAL DATA

### Multiunit activity data obtained from Utah electrode array in monkey

Now we consider data recorded from a monkey performing a visual task, where the stimulus corresponds to vertical gratings covering all recorded V1 receptive fields from the Utah arrays (see Methods for details). We aim to provide a proof of concept for the connectivity analysis for this type of data, so as to complement the more classical analysis based on the activity of individual channels; therefore we do not focus on comparing the 4 stimulus conditions with each other.

The multiunit activity envelope (MUAe) is obtained as described in Methods. In Fig. 8A, the resulting MUAe is represented for two out of the 26 channels (red and purple) for two trials in the top and middle panels, 400 ms before and 600 ms after the stimulus onset. The typical analysis of MUAe activity consists of averaging over 200 trials, which exhibits a peak immediately after the stimulus for the two channels in the bottom panel. Among the 26 channels, about a third show a large increase in activity after the stimulus onset as compared to before (namely, a post-stimulus mean activity larger by more than three standard deviations compared to the pre-stimulus activity); almost all channels show a moderate increase of one standard deviation. One channel is discarded for a much larger activity (by 5 times) than all others.

**Figure 8.**
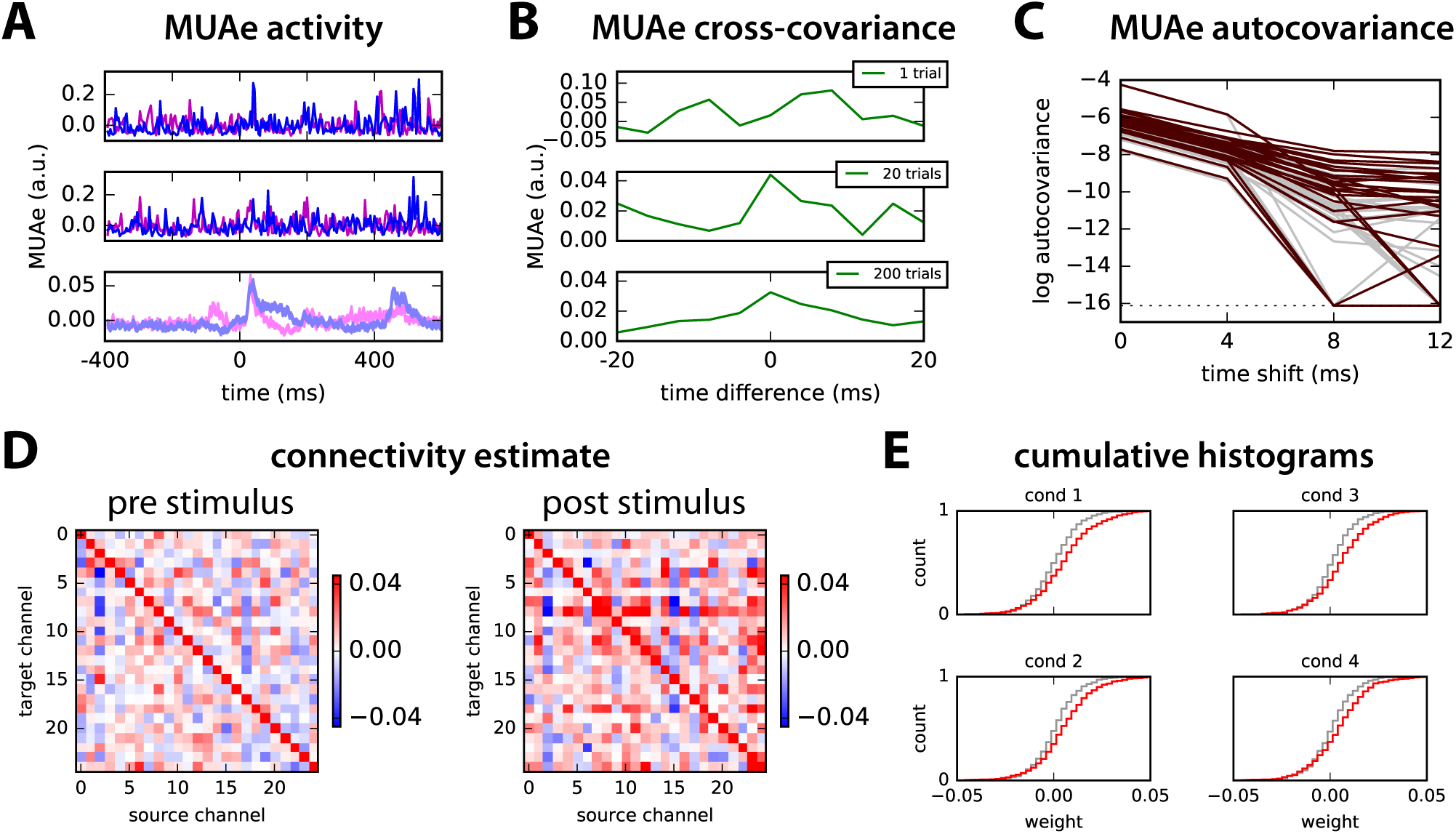
Application to multiunit activity (MUAe) data. **A:** Example of two trials (top and middle panels) of multiunit activity envelope (MUAe) for two channels of recordings using Utah electrode array in the primary visual cortex of a monkey (in arbitrary units; see text for further details). The bottom panel represents the average over 200 trials (with standard-error mean for the thickness of the curve). The stimulus is presented at time point 100 (actually 400 ms, since the smoothing corresponds to a smoothing window of 4 ms). **B:** Example of cross-covariance between the two channels in A averaged over 1, 20 and 200 trials. **C:** Autocovariance profiles of MUAe signals for all 25 channels and time shifts up to 12 ms averaged over 200 trials plotted with a log y-axis: comparison of signals before (gray) and after (red) the stimulus presentation. **D:** MVAR estimates of the connectivity between the 25 channels for the MUAe activity 200 ms before and after stimulus presentation (i.e., 50 time points each), averaged over 200 trials. The scaling has been optimized to enhance the legibility off-diagonal elements. **E:** Comparison of the cumulative distribution of connectivity weights (off-diagonal elements in D) for the 4 conditions. Gray and red indicate before and after the stimulus, respectively.

To further investigate the temporal information conveyed by MUAe jointly for pairs of channels, we calculate the pairwise covariances between them, after centering the MUAe activity individually for each trial. Fig. 8B shows the stabilization of the cross-covariance between the two channels in Fig. 8A from a single trial to averages over 20 and 200 trials. Note the asymmetry with respect to time difference: this information is extracted by the network model to estimate the interactions between the neuronal populations recorded by the channels. Then we verify that the model can be applied to these data, by examining the MUAe autocovariances in Fig. 8C, which exhibit a profile corresponding to an exponential decay up to two time shifts (i.e., 8 ms for the downsampling every 4 ms), that is, a straight line in the log plot. This suits an autoregressive model with large positive values on the diagonal of the connectivity matrix *A*.

Both connectivity matrices for the 25 channels estimated using the MVAR method before and after the stimulus are illustrated in Fig. 8D for condition 1: we find larger off-diagonal values for the period after the stimulus than before. This is actually true for all conditions, as indicated by the more spread distributions in red as compared to gray in Fig. 8E. The channels appear to be coordinated at the considered time scale of 4 ms and their collective interaction scheme is affected by the stimulus presentation.

### Significance test for real data: interactions related to stimulus presentation

We then use the local and global tests based on 1000 surrogates (with random permutation) to retain only significant interactions from the estimates in Fig. 8D: this leaves a few interactions for the pre-stimulus period in Fig. 9A (left panel), 8 out of 650, which is of the order of the desired false-alarm rate set to 1% (namely, the extreme 0.5% of each tail). In contrast, many more post-stimulus interactions survive the significance tests in the right panel: almost all these interactions are unidirectional. The counterpart for circular shift for Fig. 9B involve 24 interactions in common with Fig. 9A. On average over the 4 conditions, 22 post-stimulus interactions are common between the two shuffling methods, to be compared with 7 for the pre-stimulus period (both with a standard deviation of 4); this corresponds to 3.5% of all possible interactions. Almost all detected interactions are unidirectional, as illustrated in Fig. 9C for both local and global tests for the post-stimulus period. Varying the threshold on the tail of the null distributions, we see that the number of detected interactions is close to the desired false-alarm rate for the pre-stimulus period in Fig. 9D (dark red and black curves, respectively). In contrast, post-stimulus interactions are many more for both local and global tests (light red and gray). The global test detects fewer interactions than the local test, indicating the necessity to take into account the disparities across channels. Around 57% of post-stimulus interactions detected by the global test (largest values in absolute value) are found by the local test.

**Figure 9.**
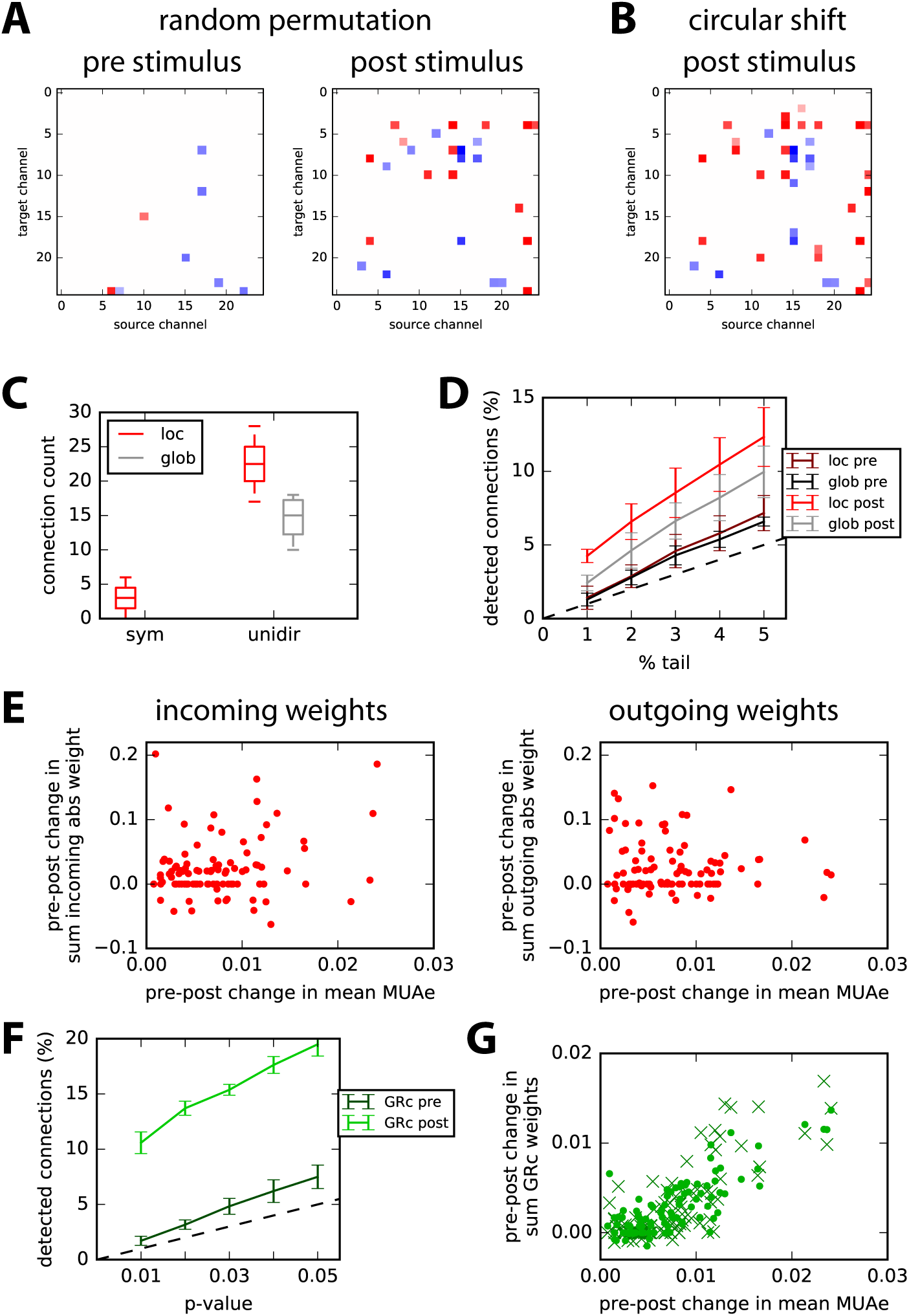
Detection of significant interactions. **A:** Examples of significant interactions with the time-rolling surrogate method; left and right panels correspond to pre- and post-stimulus periods, respectively. The p-value corresponds to the upper and lower 0.5% tails of 1000 surrogates (local test). Many more interactions are found for post- than pre-stimulus period. **B:** Matrix of post-stimulus interactions where green pixels indicate symmetric and purple asymmetric significant interactions; other are left blank. **C:** Comparison of number of asymmetric interactions versus symmetric interactions for the local (red) and global (gray) tests over the 4 conditions. **D:** Ratios of detected interactions for pre- and post-stimulus periods with the desired false-alarm rate equal to 1 to 5% corresponding to both local and global tests. **E:** Change between pre- and post-stimulus periods of the sum of incoming (left) and outgoing (right) significant weights - in absolute value - plotted against the change in the change in mean MUAe. Each dot represents a channel and all 4 conditions are grouped here. **F:** Similar plot to D for parametric GRc. **G:** Similar plot to E with sumsofGRc values for each node over its incoming (dots) and outgoing (crosses) connections. Only significant interactions passing the parametric test with p-value of 0.01 in F are retained here.

Finally, we check the relationship between the strengths of significant interactions - in absolute value - and the increase of average MUAe observed in Fig. 8A (lower panel). In Fig. 9E, the plotted dots correspond to the pre-post change in the sum of incoming (left panel) and outgoing (right panel) significant interactions for each node. The summed interaction values positively correlate with the MUAe difference (post minus pre) only for the incoming interactions: *p* = 0.03 with a coefficient of 0.21; nevertheless, the plotted values exhibit a large variability, which moderates this significance. In contrast, outgoing interactions exhibit a non-significant negative correlation (*p* ≫ 0.1). This suggests a stimulus-driven gating of the effective gain for incoming anatomical connections to recorded cell populations. The application of our method thus unravels stimulus-driven directed interactions and cannot be merely explained by an increase of single-channel MUAe activity.

In comparison, similar detection with parametric GRc testing gives more interactions for the post-stimulus period in Fig. 9F (bright green), more than twice the number for a p-value of 0.01 (corresponding to 1% in Fig. 9D). Moreover, the distributions of MVAR coefficients in Fig. 9E and the corresponding ones for GRc estimated values have similar KS distance when comparing - for each condition - the pre- and post-periods (mean of 0.17 with a std of 0.01 over the 4 conditions). This means that GRc values collectively discriminate between the two periods as well as the estimated MVAR coefficients. However, the pre-post changes in incoming and outgoing GRc values strongly correlate with the change in mean MUAe (*p* < 10^−20^ for both incoming and outgoing connections); note that the estimated connectivity is not symmetric, though. This contrasts with the two plots in Fig. 9E and suggests that the two methods may capture distinct effects at work in the network of neuronal populations. Note that the non-parametric method for GRc did not work for the experimental data here, failing to detect more interactions than the expected false-alarm rate.

## DISCUSSION

### Non-parametric MVAR-based detection of linear feedback in recurrent networks

This paper proposed a non-parametric method to detect pairwise feedback connections in biological networks with possibly strong and/or dense recurrent feedback. We examined the benefit of detecting directional connections in MVAR-like models estimated using the OLS autoregressive coefficients instead of the error residuals ratios (Fig. 1). To our understanding, the good performance of the presented method has three reasons. First, the ROC-based prediction power in Fig. 2, which relies on the estimated ranking of connections in the network (i.e., from small to strong weights), is more robust for the regression coefficients than residual log ratios for recurrent networks with relatively large density (0.1 – 0.3%); these networks overall imply many redundancy and convergence patterns of connections (M. Ding et al., 2006; Stramaglia et al., 2014). Second, practical coefficient-based connectivity detection performed using non-parametric significance tests based on time-series randomization (red distribution in Fig. 4B) yields better results than conditional Granger causality ratio test in the time domain, either with the standard parametric F test (Barnett and Seth (2014), dashed line) or with non-parametric testing (green and blue-green distributions). Finally, the use of connection-specific significance testing achieves higher accuracy than a network-pooled alternative, especially when asymptotic assumptions do not hold (e.g., small number of time samples), as illustrated by the local non-parametric test in Figs. 3F and 4B (red versus dark gray). Together, our results highlight the need for testing strategies that capture the heterogeneity of sufficiently large networks to detect individual connections. Further note that assessing the connectivity via the regression coefficients space brings an additional advantage for network studies: the estimated connection weights can be interpreted and compared across the whole network, for example using graph theory (Sporns, 2013).

Our approach for generating surrogate distributions can be encompassed in the family of constrained randomization methods (Schreiber, 1998; Schreiber & Schmitz, 1996). Here we have shown that random permutation provides a good estimation of all types of connections (Fig. 5) in false-alarm and miss rates. Comparatively, the circular-shift method performs as well except for self connections that are not detected at all; this also holds for non-random topologies (results not shown). Hence, preserving the autocovariance structure in the generation of surrogates does not provide a substantial advantage here (Fig. 5B-C). Both methods show a good control for the false-alarm rate in comparison to phase randomization (Schreiber & Schmitz, 1996) and a control Gaussian approximation over the whole network (STD), which lead to an excess of about 1% of false alarms (i.e., ~ 50 connections for a network of 70 nodes). These results show the importance of choosing a surrogate method adapted to the detection problem. For distinct dynamics governing the nodal activity such as nonlinearities, conclusions may differ and further research along these lines is necessary.

As mentioned earlier, the use of an individual null distribution for each connection (local test) gives better results for the miss rate (by a few %) than lumping together all matrix elements of all surrogates (global test), provided sufficiently many surrogates are generated. For the size of networks considered here, computation time is not an issue (Fig. 4D) and our results support the choice of the local test over the global test to attain between accuracy in the true-positive detection. This may be especially true for specific topologies or networks with both excitatory and inhibitory connections, see Fig. 6. In other words, the local test incorporates to a better extent the network heterogeneities in order to build the null distribution for each connection. The present study was limited to ordinary least-square (OLS) estimates for MVAR, but there exist alternative estimators such as the locally weighted least-square regression (Ruppert & Wand, 1994) that may perform better for particular network topologies. The extension of the presented surrogate techniques to the case where observations are sparser than connections - implying that the covariance matrix is not invertible - is another interesting direction to explore (Castelo & Roverato, 2006).

The problem of multiple comparison is intrinsic to brain connectivity detection as the number of testable connections across brain regions is massive (Rubinov & Sporns, 2010). In this context, different approaches have been developed to control the family-wise error rate in the weak sense. For instance, many studies on neuroimaging data (Genovese, Lazar, & Nichols, 2002; Nichols & Hayasaka, 2003) or electrophysiology (Lage-Castellanos, Martínez-Montes, Hernández-Cabrera, & Galán, 2010) have resorted to procedures that control the false discovery rate (FDR) (Benjamini & Hochberg, 1995), namely, the expected number of falsely declared connections among the total number of detections. These methods make decisions on single connections relying on the entire sequence of p-values computed for each connection and yield substantial statistical power gains over more conservative methods such as Šidák-Bonferroni (Abdi, 2007). With the ever growing application of graph theory to brain connectivity, new methods have been proposed that exploit the clustered structure of the the declared connections (Han, Yoo, Seo, Na, & Seong, 2013; Zalesky, Fornito, & Bullmore, 2010) to propose cluster-based statistical tests (Maris & Oostenveld, 2007) that attain similar performance to FDR methods. The present work can therefore be understood as a primary step before performing any or several multiple-correction procedures. By defining an accurate null model of inexistent connections, p-value estimates per connection are improved and cluster-based surrogate distributions can be better approximated, which is expected to empower the overall control of false positive rates in network connectivity analysis.

### Applications to real electrophysiological and neuroimaging data

A motivation for our method is the detection of neuronal interactions between electrode channels, which is often performed using spectral Granger causality analysis on local-field potential (low-passed signal of electrode measurements) or ECOG measurements. As an alternative, we have applied our connectivity detection method to MUAe recorded from macaque area V1 in order to provide a proof of concept. Multi-channel recording devices have been developed in the past years to obtain this type of data (Fan et al., 2011; Roy & Wang, 2012) and a recent cognitive study has highlighted group properties of MUAe activity for a similar experimental setup to the one used here Engel et al. (2016). Looking at the variability for individual trials in Fig. 8A (upper and middle panels), it is rather surprising that the MUAe conveys temporal information about joint activity for pairs of channels (Fig. 8B), which can be be related to causal directed interactions. This means that the high trial-to-trial variability is not an absolute limitation to temporal coordination, even though the latter only becomes apparent over multiple trial repetitions, just as the post-stimulus increase of MUAe for single channels (lower panel in Fig. 8A). The procedure detects a number of significant directional interactions well above the false positive rate that were not merely explained by the MUAe increase (Fig. 9D and E). In comparison, the control for the pre-stimulus period in the 4 different conditions detects just above the expected false-alarm rate. We find that the parametric Granger causality test also detects many more interactions after the stimulus than before; however, these interactions happen to link channels exhibiting the strongest increases in MUAe activity. This presents the caveat of providing little information in addition to the changes observed at the single-node level, when interpreting the obtained results. The research of optimal preprocessing - in particular the filtering to obtain MUAe - to obtain a robust detection of interactions is left for a later study. Likewise, such electrode recordings are often analyzed with respect to specific frequency bands (e.g., alpha and gamma), but the adaptation of our framework along the lines of previous works (Dhamala et al., 2008; L. Ding et al., 2007) and comparison of detected interactions with established methods will be done in future research.

More generally, our methodology requires adequate preprocessing of multivariate time series - activity aggregation over hundreds of voxels for fMRI and 4-ms smoothing of MUAe for electrode recordings - such that the autocovariance profiles match the exponential decay of the dynamic network model with linear feedback, which underlies the connectivity analysis. Although filtered and smoothed signals fall into the class of autoregressive moving-average (ARMA) models, our approach is applicable provided the autocovariances exhibit a profile resembling Fig. 8C. We further expect non-parametric testing methods for be extendable to ARMA processes, complementing approaches developed by Barnett and Seth (2015); Friston et al. (2014). In theory, stationarity of the time series remains a critical issue as we need sufficiently many observed samples to obtain precise covariances from which we estimate the connectivity, but this may not be a strong limitation for MUAe in practice in view of the post-stimulus average response shown in Fig. 8A.

## ACKNOWLEDGMENTS

MG acknowledges funding from the Marie Sklodowska-Curie Action (grant H2020-MSCA-656547). MG and GD were supported by the Human Brain Project (grant FP7-FET-ICT-604102 and H2020-720270 HBP SGA1). GD and ATC were supported by the European Research Council Advanced Grant DYSTRUCTURE (Grant 295129). AT was supported by the UK Medical Research Council (grant MRC G0700976). The authors are grateful to Robert Castelo and Inma Tur for constructive discussions.

## AUTHOR CONTRIBUTIONS

Project was formulated by MG and ATC. Simulation code was developed by MG. Experimental data were provided by AT and XC. Manuscript was written by MG, ATC, AT and GD.

